# Protracted development in children of perceptual segregation of competing talking faces in the multisensory cocktail party problem

**DOI:** 10.64898/2026.03.20.706527

**Authors:** Katia Steinfeld, Micah Murray, David J. Lewkowicz

**Affiliations:** The Radiology Department, Lausanne University Hospital and University of Lausanne, 1011 Lausanne, Switzerland; The Sense Innovation and Research Center, 1007 Lausanne and Sion, Switzerland; Child Study Center, Yale School of Medicine, New Haven, CT, USA

## Abstract

Successful communication with our social partners requires binding, integrating, and perceptually segregating the audible and visible attributes of the multiple talking faces that we often encounter in social situations, a challenge known as the multisensory cocktail party problem (MCPP). Although audiovisual (AV) temporal synchrony is a powerful cue for binding speech signals, how children develop the ability to use this cue to segregate a target talker remains unclear. Here, we examined the development of gaze dynamics supporting multisensory segregation in 3-7-year-old children (N = 149) and adults (N = 37) viewing four talking faces accompanied by a single auditory utterance synchronized with one of the faces (i.e., target). Using metrics of gaze dynamics from information theory, namely proportion of total looking time, stationary entropy, transition entropy, and transition rates, we show that even though sensitivity to AV synchrony is present by age 3, it is insufficient for efficient target segregation. It is not until ages 5-6, following a qualitative shift in dynamic gaze control and more structured distractor transitions, that target selection becomes more efficient, but still not as efficient as it is in adults. We interpret these developmental changes as reflecting a shift from early detection of multisensory cues to later-emerging strategies that organize visual sampling in relation to auditory information in a task-dependent manner. Together, they demonstrate that solving complex multisensory challenges depends on AV integration as well as on the development of dynamic gaze organization that supports efficient multisensory perceptual segregation over time.

**Significance Statement:** Social communication requires segregating one talker from others, a challenge known as the multisensory cocktail party problem. Although adults solve this efficiently, how this ability develops remains unclear. Using dynamic gaze measures derived from information theory, we show that multisensory segregation in childhood depends not only on detecting audiovisual synchrony but also on the emergence of structured gaze strategies. Only by ages 5-6 do children combine sustained target fixation with organized sampling of competing talkers. Even by age 7, these audiovisually guided strategies remain immature relative to adults. These findings reveal probabilistic sampling mechanisms through which gaze supports multisensory segregation, offering a mechanistic account of how children learn to navigate complex social environments, with implications for language development and education.

---

In everyday social settings, communication requires identifying one talker among other, competing talkers, a challenge known as the multisensory cocktail party problem (MCPP) (1–3). Solving the MCPP requires binding and integrating auditory and visual speech cues originating from a single talker and segregating them from those of competing talker. Because AV speech signals co-vary continuously over time, successful segregation depends on dynamic multisensory processes rather than static sensory cues.

Temporal coherence between auditory and visual speech provides a powerful cue for AV binding (4–7), rendering integrated AV objects and events perceptually more salient than their unisensory counterparts (8–10) and thus facilitating their perceptual segregation (11–14). Eye movements offer a unique window into these processes, as gaze reflects how perceivers sample, prioritize, and update information over time. Previous work has shown that eye movements track perceptual organization in vision (15, 16), prioritize relevant auditory speech features (17), and index AV speech integration in both infants (18) and adults (19, 20). Together, these findings show that gaze reflects the sampling of multisensory information during AV speech perception.

Using a paradigm specifically designed to study the MCPP, Lewkowicz et al. (2021, 2022) showed that both adults and 3-to 7-year-old children preferentially look at an audiovisually synchronized target talking face compared with competing distractors (i.e., audiovisually desynchronized talking faces), and that this preference increases with age. These findings were based on proportion of total looking time (PTLT), a widely used aggregate measure of gaze allocation. While informative, PTLT provides no insight into how gaze is dynamically structured within trials as participants exploit multisensory cues to identify the target. Understanding how gaze sampling is organized across successive fixations is critical because solving the MCPP requires accumulating information over multiple gaze samples rather than relying on static cues.

Here, we address this limitation by employing measures of gaze dynamics derived from information theory, including stationary and transition entropy (21), to eye-tracking data from adults and children who participated in Experiment 1 of the Lewkowicz et al. (2021) and (2022) studies. Our goal was to determine whether and how dynamic gaze organization changes across development and to identify the mechanisms underlying the emergence of efficient multisensory segregation. We expected that the multisensory binding and integration processes necessary for successful perceptual segregation gradually improve during early childhood. Moreover, we hypothesized that sensitivity to AV synchrony is necessary but not sufficient for efficient target segregation and, thus, that the developmental emergence of an efficient AV gaze exploration strategy is also essential to solving the MCPP.

## Methods

### Participants

We analyzed eye-tracking data from 37 adults (Experiment 1 in Lewkowicz et al., 2021) and from 177 children aged 3–7 years (Experiment 1 in Lewkowicz et al., 2022). All participants had normal or corrected-to-normal vision and hearing. Detailed inclusion criteria and demographics are provided in the Methods section of the SI Appendix. All participants and their caretakers gave their informed consent. All actors appearing in the stimuli, as in Figure 3, provided informed consent for the publication of their images. This study was approved by the IRB at Northeastern University (#14-04-12).

### Apparatus and Stimuli

Child and adult participants were seated approximately 60 cm from a remote video-based eye tracker (REDn SensoMotoric Instruments, SMI), sampling at 60 Hz. Visual stimuli were presented on a laptop computer 11 × 13-inch screen (Dell Precision M4800), while the instructions and auditory stimuli were presented through headphones.

During the practice and test trials, participants saw composite videos depicted in Figure 1. These consisted of four equally sized videos of the same female face presented in each quadrant of the screen. During each test trial, all four faces articulated the same utterance and the accompanying audible utterance was the same as the visible utterance.

**Figure 1.**
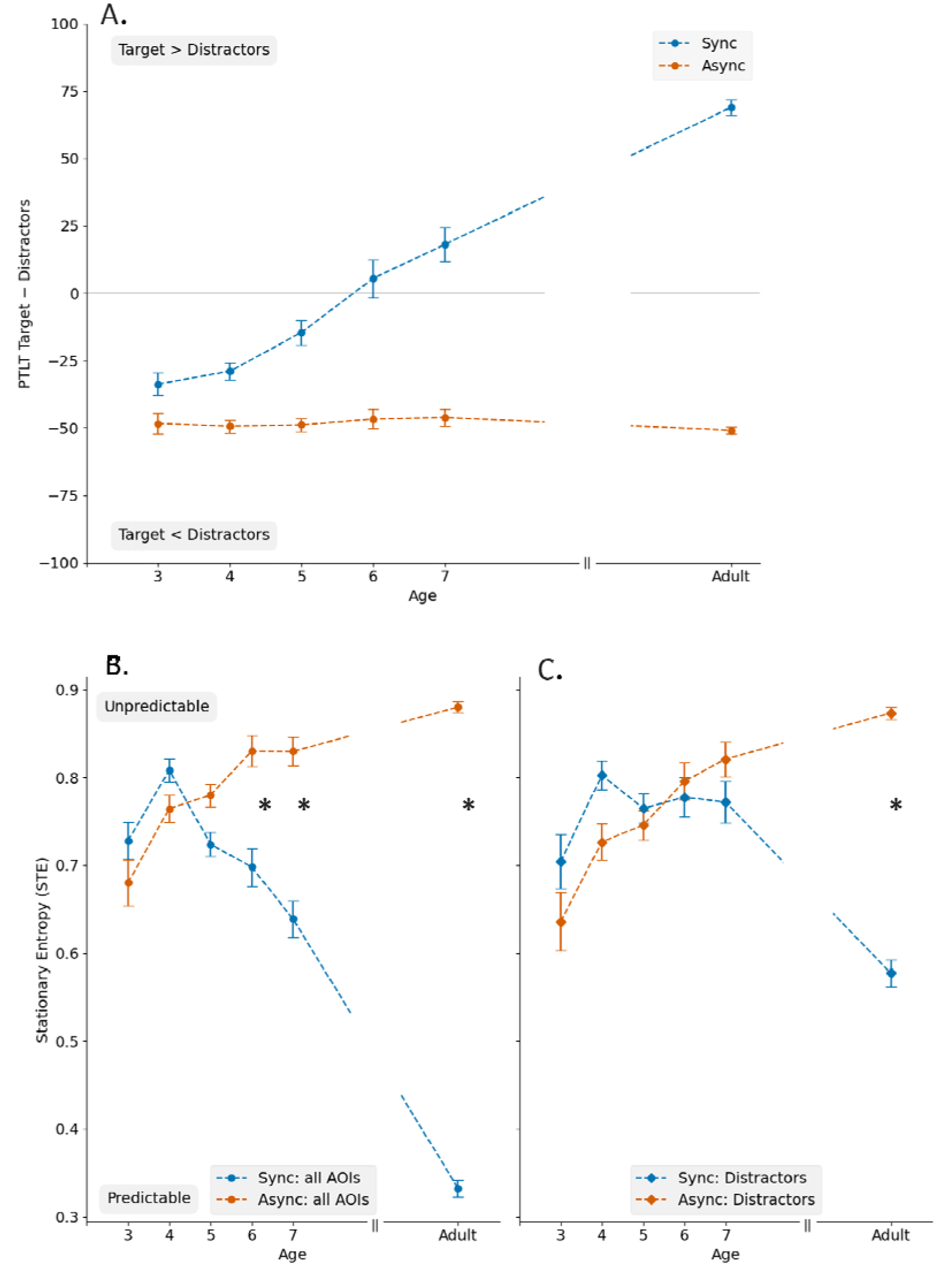
(A) Difference between target-PTLT and distractor-PTLT in Sync and Async. Stationary entropy across all face AOIs. (C) Stationary entropy across distractors. Error bars are standard errors of the mean and asterisks indicate statistically significant differences.

During half the test trials (i.e., the Sync condition), the audible utterance was temporally synchronized with the visible articulations of one of the four talking faces (the target) but desynchronized with the articulations of the other three talking faces (the distractors). During the other half of the trials (i.e., the Async condition), the audible utterance was desynchronized from the visible articulations of all four talking faces. The procedure used to create the composite videos is detailed in the Methods section of the SI Appendix.

### Procedure and Design

Measures of gaze were acquired from the right eye. A calibration phase was followed by age-appropriate instructions (see SI Appendix). There were two 15 s practice trials in which the female actor who was not the same as the actors in the test trials (see SI Appendix) was seen and heard. Then, children were given 4 pairs of trials consisting of one synchronous and one asynchronous trial according to Latin a Square design (for details see SI Appendix). Adults were given 32 test trials in one of 4 random orders. Immediately after each test trial, participants were instructed to either point (children) or press a key (adults) corresponding to the face that they thought was the talking face and were not given any feedback. Pointing responses were not recorded as they were intended as incentives.

### Data Pre-Processing and Analyses

Preprocessing and computation of dependent variables were performed in Python, and statistical analyses were conducted in SPSS. In the raw fixation data, saccades were identified using the Identification by Two Moving Windows (I2MW) (Hooge et al., 2022). No further processing of fixations was employed. Procedures and parameters are detailed in the Methods section of the SI Appendix. Fixations were labeled according to the dynamic AOI coordinates of each face in each video stimulus. Dependent variables described in Table 1 were computed over each trial (see SI Appendix for details). The corresponding hypotheses are outlined in *Supplementary Table 1* (see SI Appendix).

**Table 1.**
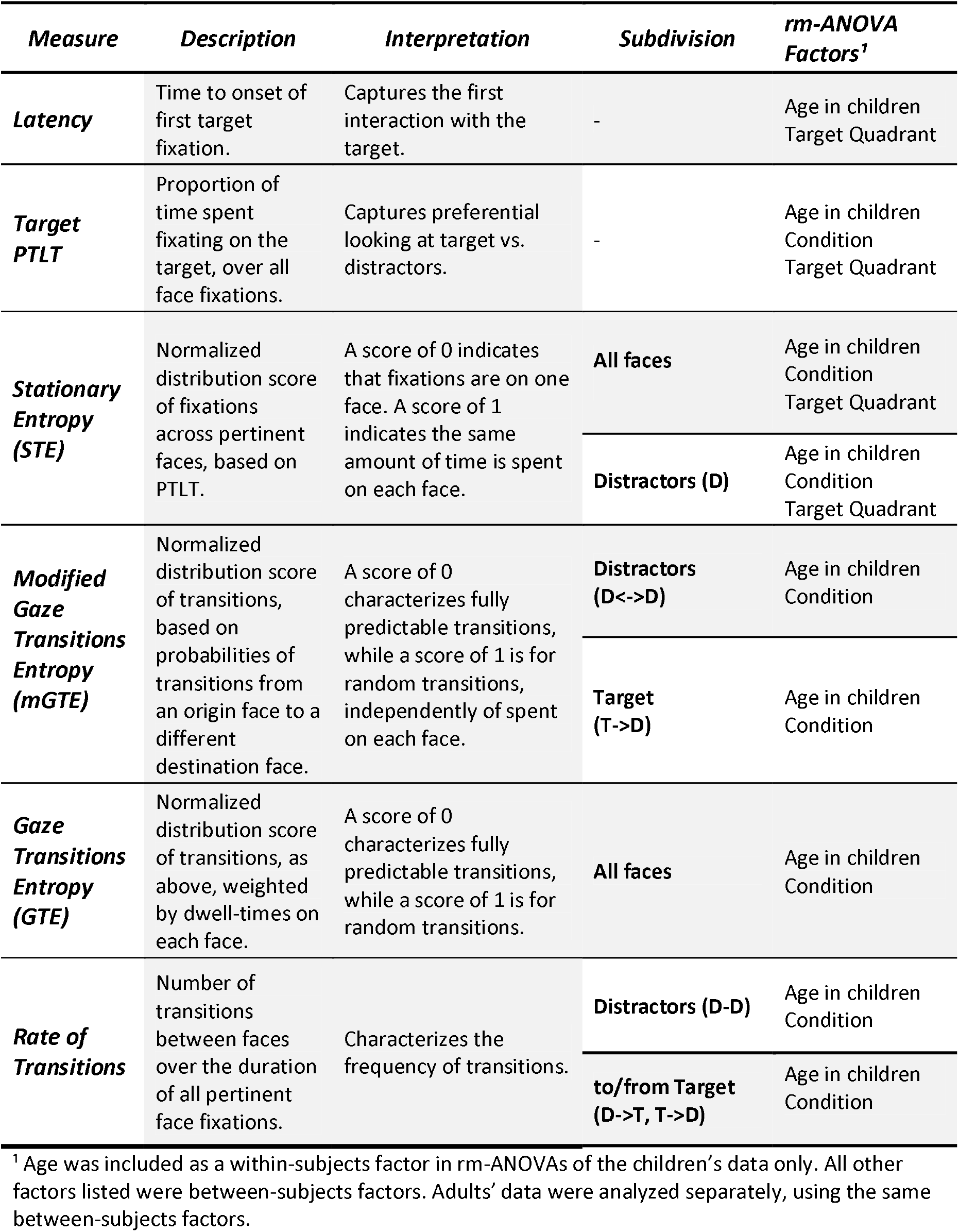
Dependent Variables Used in the Data Analyses.

Throughout the analyses, we conducted separate repeated-measures analyses of variance (rm-ANOVAs) for children and adults (see *Supplementary Tables 3-11* for the complete report of main effects and interactions). Age was included as a between-subjects factor in all rm-ANOVAs of children’s data, but not in the analysis of adults’ data (see summary in Table 1 below). Target Quadrant was included as a within-subjects factor in all analyses of fixation-based metrics (Latency, Target PTLT, STE) to investigate vertical asymmetries in dwell-time, but it was not included in the analyses of transition-based metrics (mGTE, GTE and Transition Rate) for which we had no *a priori* spatial asymmetry hypotheses (see Supplementary Table 1 in the Methods section of the SI Appendix for all hypothesis). Post-hoc comparisons were Bonferroni-corrected for multiple comparisons.

## Results

### Latency (Time to Onset of First Target Fixation)

In children, latency decreased as a function of Age (see *Supplementary Table 3*). In children and adults, latency differed across Target Quadrants, with targets in the upper quadrants detected faster than those in the lower quadrants (lower vs. upper difference mean, ± s.e.m.: children = 1.1 ± 0.2 s; adults = 0.9 ± 0.1 s).

### Target PTLT

Overall, children and adults spent significantly more time fixating the target in the Sync than in the Async condition (see *Supplementary Table 4*). In children, the significant Condition x Age interaction was due to the Sync minus Async difference reaching significance only from age 4 onward (age 3: p = .055; ages ≥4: all p’s < .001).

Target-PTLT also varied by Target Quadrant and, for children, this effect interacted with Condition and Age (see *Supplementary Table 4*). Children (all p’s < .001) and adults (all p’s < .001) fixated the upper-quadrant target longer than the lower-quadrant target. Cross-condition comparisons indicated that the Sync > Async difference was significant when the target was in the lower quadrants (all p’s<= .028) but not when it was in the upper quadrants (all p’s>= .283) at 3 years of age and that, starting at 4 years of age, children spent significantly more time fixating the target in Sync than Async regardless of its location (all p’s <= .027).

### Stationary Entropy (STE)

#### Stationary Entropy of Fixations of all Faces

Synchrony condition affected children’s and adults’ responsiveness differentially and, its effects also interacted with Age in children (see *Supplementary Table 5*). As shown in Figure 1B, from 5 years of age, STE was lower in the Sync than the Async condition (<5-year-olds, all p’s => .09, >5-year-olds all p’s <= .01) and this difference increased with age.

Stationary entropy also varied by Target Quadrant in children and adults, and this interacted with Condition in adults only (see *Supplementary Table 5*). Specifically, in children, STE was lower when the target appeared in the upper vs. the lower quadrants regardless of synchrony condition (all p’s <= .019). No other quadrant comparisons reached significance. In adults, STE was lower in Sync than in Async regardless of target quadrant (all p’s <= .001).

#### Stationary Entropy of Fixations of Distractors (D)

We observed a Condition x Age interaction in children and a significant main effect of Condition in adults (see *Supplementary Table 6*). As can be seen in Figure 1C, 3- and 4-year-olds’ STE on distractors was lower in the Async than in Sync (all p’s <= .045). This was not the case from age 5 onwards, with no difference in distractors STE across conditions (all p’s >= .094). In adults, STE on distractors was lower in the Sync than Async condition and the difference between conditions was larger than in children.

Stationary entropy also varied by Target Quadrant in children and adults, and this interacted with Condition in adults only (see *Supplementary Table 6*). In children, STE on distractors was lower when the target appeared in the upper vs. lower quadrants (all p’s <= .019) and this was independent of condition. No other quadrant comparisons reached significance. In adults, STE on distractors was lower in the Sync than the Async condition for all target quadrants (all p’s <= .001).

### Modified Gaze Transition Entropy (mGTE)

#### mGTE of transitions away from the target (T->D)

This measure varied with Age in children and as a function of synchrony condition in adults (see *Supplementary Table 7*). As can be seen in Figure 2A, mGTE away from the target was lower in 5-to 7-year-olds than in 3-year-olds (3-vs. 5-, 6- and 7-year-olds: all p’s<= .010) across conditions. No other pairwise age comparisons reached significance (all p’s>=.061). In adults, T->D mGTE was lower in the Sync than in the Async condition (T->D: p < .001).

**Figure 2.**
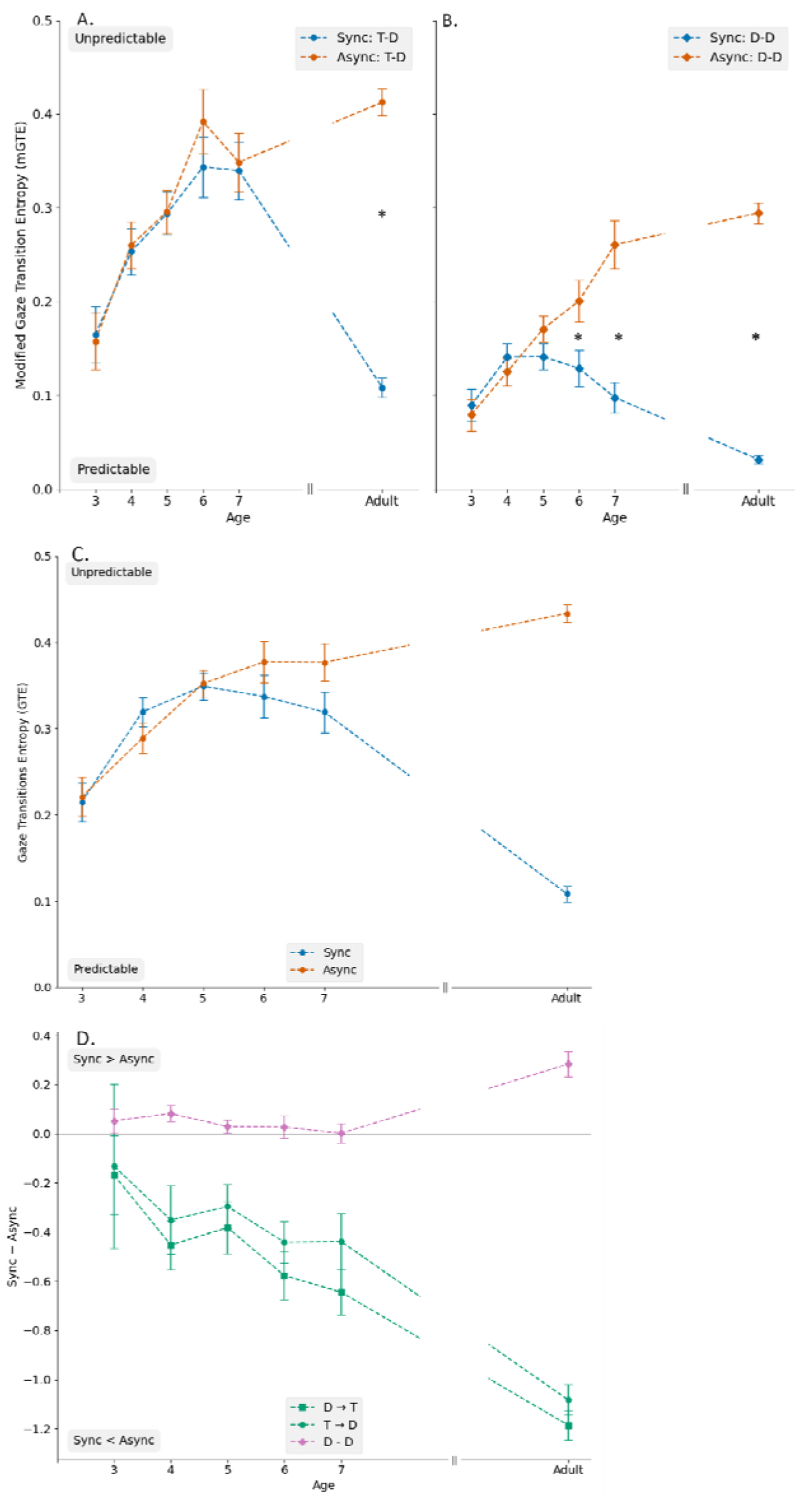
Modified Gaze Transition Entropy in Sync and Async (A) from target to distractors and (B) between distractors. (C) Gaze Transition Entropy in the Sync and Async conditions. (D) Difference between Sync and Async for rate of transitions between Target and Distractors (T->D and D->T) as well as between Distractors (D<->D). Error bars are standard errors of the mean and asterisks indicate statistically significant differences.

**Figure 3.**
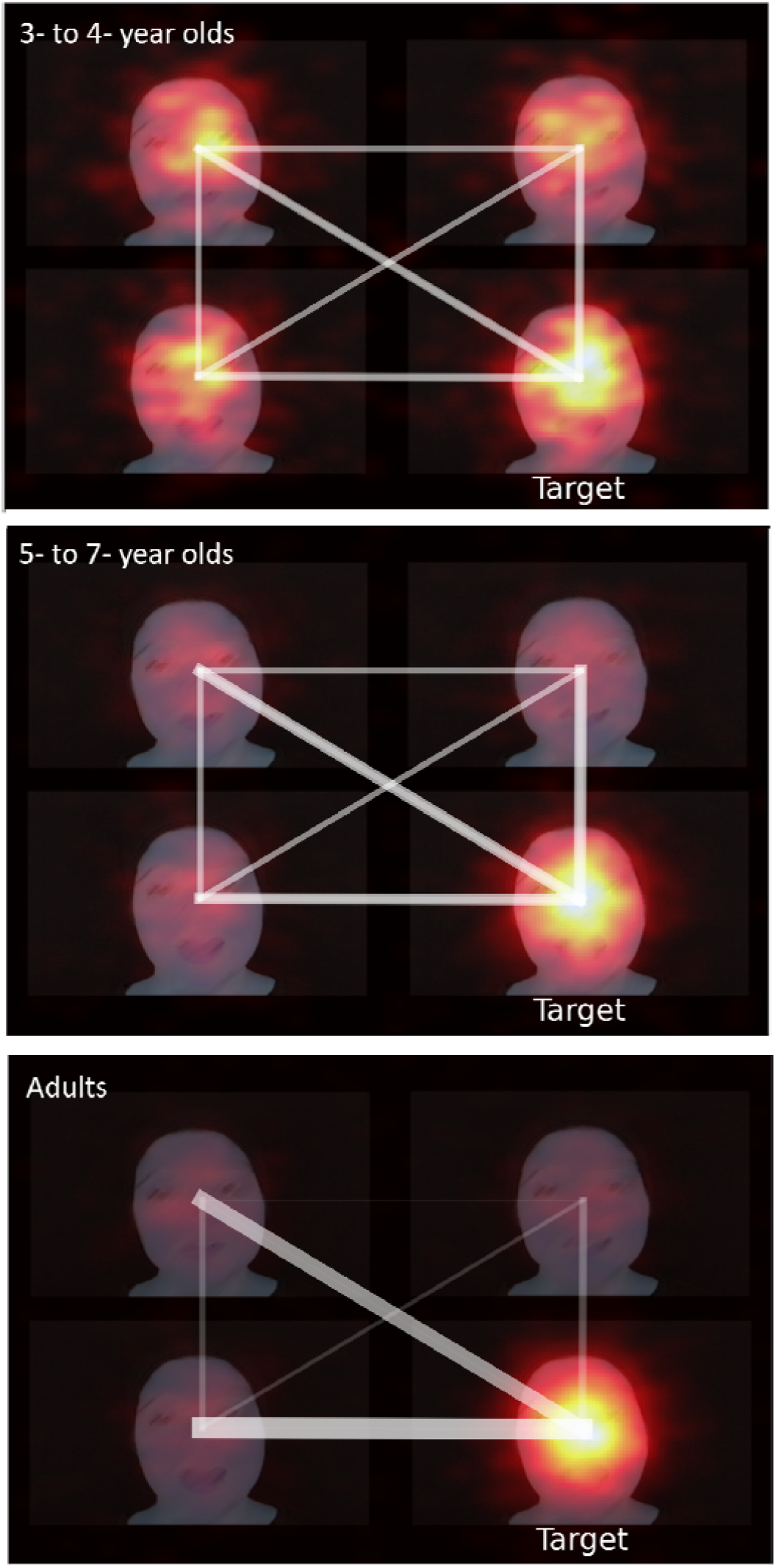
Summary cartoon depicting changes in gaze patterns in Sync vs. Async, for a lower right quadrant Target in 3-to 4-year-olds (top panel), 5-to 7-year-olds (middle panel) and Adults (bottom panel). Heatmaps represent the distribution of gaze on each AOI, with warmer colors being associated with greater PTLT and lower static entropy. Lines represent transitions between AOIs, with thicker lines being associated with lower mGTE or GTE as well as more frequent transitions.

#### mGTE of transitions between distractors (D<->D)

*D<->D* mGTE differed as a function of synchrony condition in children and adults and it also interacted with Age in children (see *Supplementary table 8*). In children (see Figure 2B) the D<->D mGTE was lower in the Sync than the Async condition from 6 years of age (Age > 6 years of age: all p’s =<.025, Age < 6 years of age: all p’s>=.187). This reflected an increase in D<->D mGTE in the Async condition as a function of age (3 vs. 5-6-7 years of age: p <= .035,, 4 vs. 7 years of age: p < .001, 5 vs. 7 years of age: p < .016; all other age comparisons: p’s >= .155) while remaining stable with age in the Sync condition (p’s >= .370). In adults, like in children from 6 years of age, D<->D mGTE was lower in Sync than Async.

### Gaze Transitions Entropy (GTE)

In children, we observed a significant main effect of Age (see *Supplementary Table 9*). As can be seen in Figure 2C, GTE was lower at 3 years of age than at all other ages (3-year-olds vs. 4-, 5-, 6-, and 7-year-olds, respectively: p’s <= .006, all other age comparisons p’s >= .157). In adults, we observed a main effect of Condition (see *Supplementary Table 8*), where GTE was lower in Sync than Async.

### Rate of Transitions

#### Rate of transitions to and from target over target fixation time

For children and adults we observed a significant Condition effect and a Direction × Age interaction for children (see *Supplementary Table 10*). As shown in Figure 2D, for children and adults transitions involving the target (T-D or D-T) were more frequent in the Async than the Sync condition (Sync minus Async < 0). D-T transitions were significantly more frequent than T-D transitions in the 3-year-olds (p = <.001) only.

#### Rate of transitions between distractors (D-D) over distractors fixation time

We observed a significant main effect of Condition in both children and adults (see *Supplementary Table 11*). The D-D transitions were more frequent in the Sync than in the Async condition and while the difference between conditions remained stable across age in the children, they were smaller than the difference in the adults (Figure 2D). Results for all rm-ANOVAs are summarized in Table 2 and represented in Figure 3.

**Table 2.**
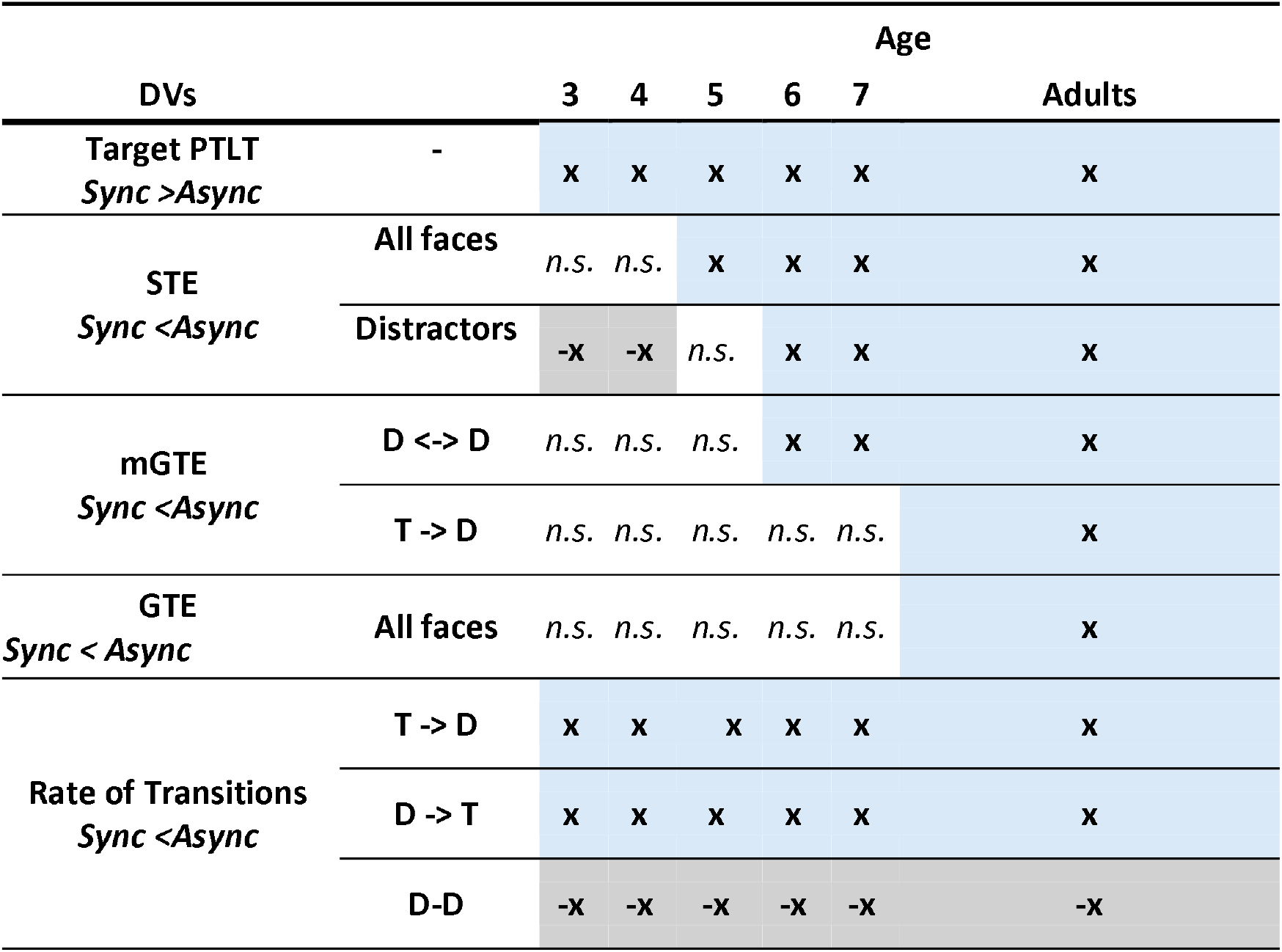
Summary of Significant Sync vs. Async Differences by Age and Dependent Measures (DVs). Cells marked with x and highlighted in blue indicate positive condition effects in the direction hypothesized and reported in the DVs column. Cells marked -x and highlighted in grey indicate a significant negative effect. Cells marked n.s. indicate non-significant effects.

## Discussion

We examined when and how the types of efficient gaze dynamics that are necessary for solving the MCPP emerge during early childhood and how they compare to those observed in adults. Although children as young as 3–4 years were sensitive to AV temporal synchrony, measures of fixation and gaze transition revealed a protracted developmental trajectory in the ability to segregate a target talking face from competing talkers. Concentration of gaze on the target increased reliably only from 5–6 years of age, while structured sampling of distractors emerged at 6 years of age and further increased into adulthood. Notably, differences between synchrony conditions in transitions away from the target were observed only in adults. Together, these findings show that early sensitivity to AV temporal synchrony is not sufficient for efficient target segregation and that the gradual emergence of dynamic gaze organization is also a necessary precondition for solving the MCPP.

### Multisensory integration at 3 and 4 years of age is not sufficient for target segregation

Using target-PTLT measures, Lewkowicz et al. (2022) found that 3- and 4-year-old children relied on AV temporal synchrony cues to identify the target talking face. In contrast, our entropy measures showed that sensitivity to AV temporal synchrony in this age range did not translate into successful fixation of the target over distractors. Instead, fixation patterns at 3 and 4 years old remained broadly distributed as shown by stationary entropy results, with no evidence of structured sampling dynamics. Selective concentration of gaze on the target talking face only emerged at 5 years of age.

The findings from the 3- and 4-year-olds are consistent with those from studies of responsiveness to AV temporal synchrony in infants (6, 7, 23–25) and from studies of synchrony-based lip reading in infancy (26, 27). Together, the evidence from infants and children demonstrates that temporal synchrony plays a major role as a multisensory binding and integration cue in early childhood (10, 24, 26–30). Importantly, however, our results show that an early-emerging sensitivity to AV temporal synchrony is not sufficient for effective perceptual segregation of a multi-talker scene. Indeed, the shift in gaze behavior observed at 5 years of age can be understood as reflecting both the emergence of multisensory binding and integration as well as the ability to flexibly weigh perceptual salience cues in accordance with task demands.

#### Spatial biases in gaze sampling constrain target segregation at 3 to 4 years of age

The developmental shift in the ability to flexibly weigh salience cues is further revealed by the spatial analysis results. Thus, at 3 years of age, a strong upper vertical bias inflated target-PTLT, resulting in reduced stationary entropy even in the absence of AV synchrony cues. By 4 years of age, AV synchrony cues allowed children to overcome this spatial bias and led to a significant increase in target-PTLT and a decrease in stationary entropy for the synchronous target at all locations. Interestingly, such spatial biases in preschoolers’ gaze allocation have been previously reported in free-viewing of complex scenes (31, 32) and natural viewing behavior (33, 34). Our results add to this evidence and support the Linka et al. (2023, 2025) view that local, low-level features and spatial biases wane in their importance for gaze behavior as development progresses. Critically, we show that before the age of 5, gaze behavior in the context of the MCPP is constrained by spatial biases to such an extent that multisensory cues, though powerful for audiovisual integration from infancy, are not yet sufficient to organize gaze sampling in accordance with the demands of this socially-relevant task.

#### The continued development of multisensory integration from 5 years of age supports sustained target fixation

Concentration of gaze on the synchronized target increased, as reflected by target PTLT and stationary entropy, between 5 and 7 years of age. Crucially, however, it did not reach adult levels. This is consistent with evidence that, in general, multisensory integration develops gradually into late childhood (35–37) and that, in particular, children’s performance in an AV integration speech task improves between early childhood and adolescence (38, 39). The protracted developmental timeline of increasing target dwell-times observed here closely parallels the types of changes in multisensory integration reported in prior studies.

#### The emergence of structured distractor sampling marks a developmental transition at 6 years of age

Our results point to a qualitative shift in gaze dynamics at 6 years of age, marked by increasingly structured sampling of distractors even though they were visually and audiovisually identical. Efficient sampling of the distractors did not depend on discrimination among them but on mapping and updating uncertainty over possible locations as evidence for the target accumulated. Thus, effective segregation required structuring exploration probabilistically, according to the remaining uncertainty about the target’s location. In adults, gaze behavior is shaped by task demands that reduce the influence of sensory salience (11, 40–43). Computational models of visual search similarly show that optimal performance depends on representing the distribution of distractors to enhance target over distractor signals (11, 44–46). This closely resembles the perceptual segregation process used for solving the MCPP, where signals from the target’s multisensory features are progressively upweighted against signals from the distractors’ features.

Transition metrics provide a dynamic behavioral signature of this segregation process. Consistent with greater dwell-time on the target, both children and adults increased their overall rate of distractor–distractor transitions when AV cues were synchronous. Despite this, however, it was not until 6 years of age that AV synchrony leads to more predictable distractor sampling (i.e., lower D<->D mGTE), although still not as pronounced as in adults. Interestingly, and in contrast, predictability of departures from the target (T→D mGTE) did not differ across the synchrony conditions at any age in children and was only evident in adults. These results reveal that a key developmental transition, marked by the emergence of structured distractor sampling, occurs at 6 years of age. This age-related pattern suggests that the development of multisensory target segregation depends on the detection of synchrony as well as the ability to update a probabilistic representation of the scene based on accumulating evidence of AV temporal synchrony.

#### Dynamic gaze transitions reveal probabilistic segregation mechanisms beyond preferential looking

Theoretically, fixation and transition measures can be framed in terms of a Bayesian approach in which participants are assumed to sample the stimulus array to assess the consistency and reliability of AV temporal synchrony cues embedded in the MCPP task. In the Sync condition, evidence can be thought of as accumulating, until posterior probabilities reach a peak and exceed a putative decision criterion. Thus, differences in dwell-time metrics (PTLT, STE) across the two synchrony conditions are related to differences in the amplitude of these posterior peaks (which are directly attributable to multisensory integration). At any one moment, the posterior distribution defines the relative probability of selecting the next face to sample, such that variations in posterior shape over time are mapped onto gaze transitions. As a result, transition entropy and transition rates represent indices of the stability and efficiency of evidence accumulation over time.

Crucially and most importantly, our gaze transition measures provide direct evidence of the development of perceptual segregation dynamics. Optimization of gaze sampling under uncertainty has previously been explored with ideal observer models. For instance, in visual search tasks designed to maximize accuracy, an ideal observer tends to fixate regions with higher uncertainty (47) and the informativeness of each fixation made by an adult human observer resembles that of an ideal observer (48). In addition, adults retain metacognitive information about previously sampled objects, allowing them to continuously assess the reliability of accumulated evidence (49). Consistent with this framework, we observed that AV temporal synchrony only coincided with more predictable departures from the target, faster and more regular sampling of distractors, and more efficient transitions across all AOIs in adults. This suggests that the ability to dynamically segregate a multisensory target embedded in a multi-talker scene, via gaze transitions under uncertainty, continues to develop beyond 7 years of age. Overall, our findings provide novel insights into the multisensory perceptual mechanisms underlying the development of social communication in early childhood.

## Acknowledgments

This work was done in part under the Multisensory Environments to study Longitudinal Development (MELD) consortium (https://lab.vanderbilt.edu/meld/), which is supported by an unrestricted gift from Reality Labs Research, a division of Meta.

## Data Availability Statement

Experimental data and analytic code will be made publicly available via OSF. Audiovisual stimulus materials contain identifiable images of human actors and are not publicly available but may be accessed upon reasonable request, in accordance with the consent provided by the actors, via Harvard Dataverse.

## References

1. D. J. Lewkowicz, M. Schmuckler, V. Agrawal, The multisensory cocktail party problem in children: Synchrony-based segregation of multiple talking faces improves in early childhood. Cognition 228, 105226 (2022).

2. D. J. Lewkowicz, M. Schmuckler, V. Agrawal, The multisensory cocktail party problem in adults: Perceptual segregation of talking faces on the basis of audiovisual temporal synchrony. Cognition 214, 104743 (2021).

3. E. C. Cherry, Some Experiments on the Recognition of Speech, with One and with Two Ears. J. Acoust. Soc. Am. 25, 975–979 (1953).

4. M. Aller, U. Noppeney, To integrate or not to integrate: Temporal dynamics of hierarchical Bayesian causal inference. PLoS Biol 17, e3000210 (2019).

5. D. J. Lewkowicz, “Development of Multisensory Temporal Perception” in The Neural Bases of Multisensory Processes, Frontiers in Neuroscience., M. M. Murray, M. T. Wallace, Eds. (CRC Press/Taylor & Francis, 2012).

6. D. J. Lewkowicz, Perception of auditory–visual temporal synchrony in human infants. Journal of Experimental Psychology: Human Perception and Performance 22, 1094–1106 (1996).

7. D. J. Lewkowicz, N. J. Minar, A. H. Tift, M. Brandon, Perception of the Multisensory Coherence of Fluent Audiovisual Speech in Infancy: Its Emergence & the Role of Experience. J Exp Child Psychol 0, 147–162 (2015).

8. P. J. Matusz, M. Eimer, Multisensory enhancement of attentional capture in visual search. Psychon Bull Rev 18, 904 (2011).

9. W. H. Sumby, I. Pollack, Visual contribution to speech intelligibility in noise. Journal of the Acoustical Society of America 26, 212–215 (1954).

10. Q. Summerfield, Lipreading and audio-visual speech perception. Philos Trans R Soc Lond B Biol Sci 335, 71–78 (1992).

11. W. Einhäuser, U. Rutishauser, C. Koch, Task-demands can immediately reverse the effects of sensory-driven saliency in complex visual stimuli. J Vis 8, 2. 1–19 (2008).

12. L. Itti, C. Koch, A saliency-based search mechanism for overt and covert shifts of visual attention. Vision Res 40, 1489–1506 (2000).

13. E. Van der Burg, C. N. L. Olivers, A. W. Bronkhorst, J. Theeuwes, Pip and pop: Nonspatial auditory signals improve spatial visual search. Journal of Experimental Psychology: Human Perception and Performance 34, 1053–1065 (2008).

14. Q. Zhao, C. Koch, Learning a saliency map using fixated locations in natural scenes. J Vis 11, 9 (2011).

15. W. Einhäuser, M. Spain, P. Perona, Objects predict fixations better than early saliency. Journal of Vision 8, 18 (2008).

16. J. M. Wolfe, Guided Search 6.0: An updated model of visual search. Psychon Bull Rev 28, 1060–1092 (2021).

17. Q. Gehmacher, et al., Eye movements track prioritized auditory features in selective attention to natural speech. Nat Commun 15, 3692 (2024).

18. I. Lozano, R. Campos, M. Belinchón, Sensitivity to temporal synchrony in audiovisual speech and language development in infants with an elevated likelihood of autism: A developmental review. Infant Behav Dev 78, 102026 (2025).

19. F. Ahmed, A. R. Nidiffer, E. C. Lalor, The effect of gaze on EEG measures of multisensory integration in a cocktail party scenario. bioRxiv 2023.08.23.554451 (2023). 10.1101/2023.08.23.554451.

20. G. Mazzi, et al., Prior expectations guide multisensory integration during face-to-face communication. PLoS Comput Biol 21, e1013468 (2025).

21. B. Shiferaw, L. Downey, D. Crewther, A review of gaze entropy as a measure of visual scanning efficiency. Neurosci Biobehav Rev 96, 353–366 (2019).

22. I. T. C. Hooge, D. C. Niehorster, M. Nyström, R. Andersson, R. S. Hessels, Fixation classification: how to merge and select fixation candidates. Behav Res 54, 2765–2776 (2022).

23. D. J. Lewkowicz, Infants’ response to temporally based intersensory equivalence: The effect of synchronous sounds on visual preferences for moving stimuli. Infant Behavior and Development 15, 297–324 (1992).

24. D. J. Lewkowicz, The development of intersensory temporal perception: an epigenetic systems/limitations view. Psychol Bull 126, 281–308 (2000).

25. D. J. Lewkowicz, Infant perception of audio-visual speech synchrony. Dev Psychol 46, 66–77 (2010).

26. A. Hillairet de Boisferon, A. H. Tift, N. J. Minar, D. J. Lewkowicz, Selective attention to a talker’s mouth in infancy: role of audiovisual temporal synchrony and linguistic experience. Developmental Science 20, e12381 (2017).

27. D. J. Lewkowicz, A. M. Hansen-Tift, Infants deploy selective attention to the mouth of a talking face when learning speech. Proceedings of the National Academy of Sciences 109, 1431–1436 (2012).

28. A. MacLeod, Q. Summerfield, Quantifying the contribution of vision to speech perception in noise. Br J Audiol 21, 131–141 (1987).

29. C. Spence, S. Squire, Multisensory Integration: Maintaining the Perception of Synchrony. Current Biology 13, R519–R521 (2003).

30. J. Vroomen, M. Keetels, Perception of intersensory synchrony: A tutorial review. Attention, Perception, & Psychophysics 72, 871–884 (2010).

31. M. Linka, H. Karimpur, B. de Haas, Protracted development of gaze behaviour. Nat Hum Behav 9, 1887–1897 (2025).

32. M. Linka, Ö. Sensoy, H. Karimpur, G. Schwarzer, B. de Haas, Free viewing biases for complex scenes in preschoolers and adults. Sci Rep 13, 11803 (2023).

33. A. Açık, A. Sarwary, R. Schultze-Kraft, S. Onat, P. König, Developmental Changes in Natural Viewing Behavior: Bottom-Up and Top-Down Differences between Children, Young Adults and Older Adults. Front. Psychol. 1 (2010).

34. O. Krishna, A. Helo, P. Rämä, K. Aizawa, Gaze distribution analysis and saliency prediction across age groups. PLoS One 13, e0193149 (2018).

35. A. B. Brandwein, et al., The development of audiovisual multisensory integration across childhood and early adolescence: a high-density electrical mapping study. Cereb. Cortex 21, 1042–1055 (2011).

36. D. Burr, M. Gori, “Multisensory Integration Develops Late in Humans” in The Neural Bases of Multisensory Processes, Frontiers in Neuroscience., M. M. Murray, M. T. Wallace, Eds. (CRC Press/Taylor & Francis, 2012).

37. M. Nardini, T. Dekker, K. Petrini, Crossmodal Integration: A Glimpse into the Development of Sensory Remapping. Current Biology 24, R532–R534 (2014).

38. R. J. Hirst, J. E. Stacey, L. Cragg, P. C. Stacey, H. A. Allen, The threshold for the McGurk effect in audio-visual noise decreases with development. Sci Rep 8, 12372 (2018).

39. S. Rohlf, L. Li, P. Bruns, B. Röder, Multisensory Integration Develops Prior to Crossmodal Recalibration. Current Biology 30, 1726-1732.e7 (2020).

40. A. Goettker, N. Borgerding, L. Leeske, K. R. Gegenfurtner, Cues for predictive eye movements in naturalistic scenes. J Vis 23, 12 (2023).

41. K. Koehler, F. Guo, S. Zhang, M. P. Eckstein, What do saliency models predict? J Vis 14, 14 (2014).

42. E. E. Oor, T. R. Stanford, E. Salinas, Stimulus salience conflicts and colludes with endogenous goals during urgent choices. iScience 26, 106253 (2023).

43. B. M. ‘t Hart, H. C. E. F. Schmidt, C. Roth, W. Einhäuser, Fixations on objects in natural scenes: dissociating importance from salience. Front Psychol 4, 455 (2013).

44. J. Gottlieb, Understanding active sampling strategies: empirical approaches and implications for attention and decision research. Cortex 102, 150–160 (2018).

45. V. Navalpakkam, L. Itti, Modeling the influence of task on attention. Vision Res 45, 205–231 (2005).

46. V. Navalpakkam, L. Itti, Optimal cue selection strategy in Advances in Neural Information Processing Systems, (MIT Press, 2005).

47. J. Najemnik, W. S. Geisler, Optimal eye movement strategies in visual search. Nature 434, 387–391 (2005).

48. S. C.-H. Yang, M. Lengyel, D. M. Wolpert, Active sensing in the categorization of visual patterns. eLife 5, e12215 (2016).

49. E. E. M. Stewart, C. J. H. Ludwig, A. C. Schütz, Humans represent the precision and utility of information acquired across fixations. Sci Rep 12, 2411 (2022).

