## Supplementary Information for "Protracted development in children of perceptual segregation of competing talking faces in the multisensory cocktail party problem"

David J. Lewkowicz

**This PDF file includes:**

Supporting Methods and Results text

Tables S1 to S11

SI References

**Other supporting materials for this manuscript include the following:**

Movies S1 to S32

Supporting Information Text

**Methods**

**Participants.** *Children*. A total of 177 children, comprising separate groups of 3- (n=23), 4- (n=41), 5- (n=55), 6- (n=27), and 7-year-old (n=31) children, contributed data in this study. Data from 5 participants were excluded because they did not complete all 8 experimental trials. Therefore, the analyses reported here were based on data from 173 children. For all comparisons involving Target Quadrants, however, data from an additional 29 participants were excluded (ten 3-year-olds; four 4-year-olds; twelve 5-year-olds; two 6-year-old; and two 7-year-olds) because they did not fixate the target stimulus in one or more Sync trials (i.e., test trials in which one of the 4 talking faces was temporally synchronized with the audible utterance).

*Adults*. A total of 38 adults, who completed all experimental trials, contributed data in this study. For the analysis of response latency as well as the effects of phase, data from 1 participant were excluded due to a failure to fixate the target stimulus in one Sync trial. As a result, this analysis was based on data from 37 adults.

**Apparatus and Stimuli**

*Procedure to Create the Composite Videos*. A crucial point to note with respect to the desynchronization of fluent audiovisual speech is that it leads to the disruption of the zero-lag temporal correlation between the continuous dynamic variations in a talker’s visible mouth movements and the accompanying audible vocalizations. This means that the temporal desynchronization of fluent audiovisual speech results in multiple points of audiovisual discordance between the visual and auditory speech streams. Also, another crucial point to note is that the movements of the vocal tract normally precede phonation by 100 to 300 ms (1) and, as a result, the onset of the auditory component of a synchronized AV speech utterance is normally delayed relative to the onset of the visual component of the utterance. This means that the visual and auditory speech streams of a fluent AV speech stream must be temporally separated by more than 300 ms if they are to be perceived as desynchronized. Accordingly, to ensure that participants perceived audiovisual desynchronization, the visual and auditory speech streams for each distractor talking face were jittered by more than 300 ms with respect to one another.

To produce the composite videos, we filmed each of the two test-phase female actors uttering two different sets of two different utterances.^[1]^ This resulted in four different videos representing the four different actor/utterance combinations. We then used each of these four videos to construct four pairs of test trials. Each pair consisted of a synchrony and an asynchrony test trial (Movies S1 to S32). The two types of test trials that made up each pair were the same except that the utterance spoken by the target face was temporally synchronized with the auditory utterance in the synchrony test trial but desynchronized with it in the asynchrony test trial. This made it possible to directly compare selective attention to a particular talking face in a particular quadrant as a function of synchrony condition.

**Procedure and Design**

*Instructions.* The instructions were as follows: “Today, we are going to play a game. The game will only continue if you are looking at the screen. If you look away, it will stop. You will see four people talking but you will only hear one person talking. Watch carefully and see if you can match the face to the voice. Then, point to the face that you thought was talking. Sometimes it will be easy and sometimes hard. Watch carefully!” During these instructions, children saw a still and silent image of four identical faces. For adults, the instructions were to indicate “the talking face” by pressing one of the keys on the numerical keypad that corresponded spatially to that talking face.

*Practice Trials.* During the first practice trial, they saw the four faces articulating the same utterance and they heard a temporally synchronized audible version of the same utterance with one of the talking faces. During the second practice trial, participants saw the same four talking faces again except that here the auditory utterance was desynchronized from all four talking faces. After each practice trial, participants saw a composite video of the four faces seen in the previous composite video except that now the faces were still and blinked occasionally. Children were instructed to point to the face that matched the voice whereas adults were instructed to press a key on the keyboard corresponding to the face that they perceived as the talking face in the preceding composite video. No feedback regarding accuracy was given after the participants pointed. Once the practice trials were completed, the participants were asked if they had any questions and, if they had none, the experiment continued.

*Latin-Squares Design for Children.* The Quadrant groups differed in terms of the location of target-face presentation for each unique pair of synchrony/asynchrony test trials, with two constraints. The first was that, within each group, the quadrant of target-face presentation for each pair of synchrony/asynchrony test trials had to be counterbalanced across the four quadrants. The second constraint was that, across the four groups, the specific quadrant in which the targets in a given pair of test stimuli were presented had to be counterbalanced. Assignment to one of the four Quadrant groups was randomly determined. For the adults, the 32 test trials were presented in one of 4 randomly generated test trial orders where, in contrast to children, synchronous and asynchronous trials were not presented in pairs. The adult participants were randomly assigned to one of the 4 orders.

**Data Pre-Processing and Analyses**

*Preprocessing.* The I2MW algorithm consists of two rolling windows which track non-linearities in Euclidian coordinates. The windows were separated by one sample and each window computes the median of the gaze position in Euclidian coordinates. If the difference between the medians computed by each window exceeds a 0.1 deg threshold, the sample between the window is considered a saccade. The labeling of saccades was manually verified, trial-by-trial, with no changes made to the output of the algorithm. The remaining traces were labeled as fixations.

*Dependent Variables: Latency*. For each trial in the Sync condition, the first fixation on the target face AOI was identified and the duration from the onset of the trial to the onset of this fixation defined the latency of the response time to the target face. In the Async condition, response latency was measured to the virtual target face, namely the face in the same quadrant where the target face was presented during the Sync trial.

*Dependent Variables: Target PTLT*. We used the same method as Lewkowicz et al. (2021; 2022) to calculate the PTLT scores for looking at the target but a different method to calculate the scores for looking at the distractors (for calculations identical to those used by Lewkowicz et al., (2021; 2022) see *Supplementary* *Table 1)*. Here, to calculate the PTLT for the target, we divided the total amount of looking at the target face AOI by the total amount of looking at the four face AOIs for each participant and trial. To calculate the PTLT for the distractors, we summed the total amount of looking time at the distractors and then divided by the total amount of looking time at the four face AOIs for each participant and trial. As a result, the Distractors PTLT was equivalent to formula 1 – target PTLT. Given that our method for calculating the Distractors PTLT resulted in this measure being a complement to Target PTLT, we did not include it in the repeated-measures (rm)-ANOVA.

*Dependent Variables: Stationary Entropy (STE)*. To characterize the spatial distribution of fixations across the 4 talking faces, we computed stationary entropy as described in Shiferaw et al. (2019). We did so both across all faces and across distractors only, where for each relevant face *i*, we computed the probability of fixation $p_{i}$ as the total duration of fixation on *i* divided by the total duration of fixations on all faces or on distractors respectively.

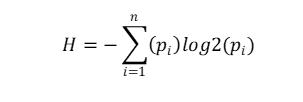

We then normalized the entropy (*H)* by the maximum possible entropy for a system with $n$states and no self-transitions, where $n$ = 4 for all faces and $n$ *= 3* for distractors, as $\log_{2}(n-1)$. It yielded values between 0 (minimal entropy), when all fixation time was on one face, and 1 (maximal entropy), when equal amounts of time were spent on all relevant faces.

*Dependent Variables: Modified Gaze Transitions Entropy (mGTE).* Because participants were tasked with identifying a target among multiple competing distractors, we sought to explore the structure of gaze transitions from target to distractors (T->D) and between distractors (D-D), independently of the stationary gaze distribution. We thus computed a variant of gaze transition entropy in which only the transitions between the relevant subset of AOIs (T->D or D-D) were considered. In doing so we did not assess the overall gaze transition complexity (see classical GTE below) but instead asked how predictable were gaze transitions that originated from the target face or from distractor faces, regardless of how often each face was visited. For each trial we removed self-transitions and constructed a 4x4 transition count matrix $C_{ij}$. Each row was normalized $p(i,j)$, with values ranging from 0 to 1, representing the probability of transitioning to a destination face *j* given an origin face. We calculated this measure separately for transitions involving the target and the distractors (T-D) and transitions among distractors only (D-D). For T-D transitions, all four faces were retained in the matrix, but only transitions in which either the source or destination was the target were counted. Similarly, for D-D transitions, only transitions in which the target was not present were counted. We then calculated row-wise conditional entropy, $H_{i}$.

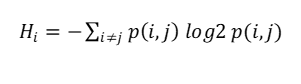

Rows with no outgoing transitions and rows with only one possible outgoing transition contributed 0 bits. For T-D entropy, this meant that only transitions originating from the target could contribute more than 0 bits, effectively representing the predictability of transitions departing from the target (T->D). For D-D entropy, this meant that transitions originating from any of the three distractors could contribute more than 0 bits, representing the predictability of all D<->D transitions. The transition entropy (*H*) for the subset (*S)* (T->D or D<->D) in a given trial was then defined as the weighted mean of the row entropies ( $H_{i})$ using as uniform weights 1/ $k$ , where *k* was the number of possible origin AOIs ( *k*=1 for T-D, *k*=3 for D-D).

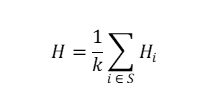

We report this quantity normalized by log2(*k*-1), which equals the maximum possible entropy between *k* states when self-transitions are excluded (log2(2) for T-D and log2(3) for D-D), to obtain values on a 0–1 scale. Effectively, the T-D transition entropy here represents how much structure is in the transitions from the target to the distractor, independently of dwell times on target. In turn, D-D transitions represent how represents how much structure is in the transitions between distractors, independently of dwell-times on each distractor.

*Dependent Variables: Gaze Transition Entropy (GTE).* To quantify how gaze dynamics varied as a combination of the spatial distribution of fixations and the structure of transitions, we computed transition entropy as in Shiferaw et al. (2019) across all faces. For each trial we removed self-transitions, so that only transitions between faces remained. Row-wise entropy was calculated as above, yielding a 4x1 vector $H_{i}$.

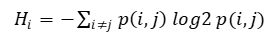

Rows with no outgoing transitions contributed 0 bits. The transition entropy for the trial was then defined as the weighted mean of the row entropies, where weights were dwell-times on each origin face divided by total dwell-time on all faces – i.e. the probability of fixation $p_{i}$ on each AOI, in a 4x1 vector. We report this quantity normalized by log2(3) to obtain a 0–1 scale.

*Dependent Variables: Rate of transitions*. Transitions were defined as consecutive fixations on different faces. Transitions between faces and non-AOI coordinates and consecutive fixations on the same face were excluded from this count. We calculated this measure for transitions involving the target and a distractor (T-D), and transitions among distractors only (D-D). Counts of transitions between distractors were not directional and were divided by the total duration of fixations on distractors during the trial. In contrast, counts of transitions between target and distractors were directional, differentiating between transitions from target to distractors (T-D) and transitions from distractors to target (D-T), and divided by the total duration of fixation on the target. Dependent variables and their associated hypothesis are listed in Supplementary Table 1.

**Results**

*PTLT as calculated Lewkowicz et al. (2021, 2022), where Distractor PTLT corresponds to the average PTLT of all distractors.* Comparisons between Stimulus Type (Target or Distractors) for each Age and Condition confirmed that, as reported in Lewkowicz et al. (2021, 2022), children from 3y.o. and adults selectively attended more to the target than to the distractors in the Sync condition (all ps=<.01), but not in the Async condition (all ps>= .215) (see Table S2). In Sync, the difference between Target and Distractors PTLT increased from 3-year-olds (mean ± s.e.m Target: 33.1±3.1%, Distractors: 22.3±1.8%) to 7-year-olds (mean ± s.e.m Target: 59±2.7%, Distractor: 13.6±.9%), without however reaching adult levels (mean ± s.e.m Target PTLT: 77.7± .8% , Distractor PTLT: 4.35± .2%).

Tables

Table S1. Dependent variables and their corresponding working hypothesis.

| **Measure** | **Subivisions** | **Hypothesis** |
| --- | --- | --- |
| **Latency** |  | Decreases with age |
| **Target PTLT** |  | Sync > Async, Sync-Async difference increases with age |
| **Stationary Entropy (STE)** | **All faces** | Async > Sync, Async - Sync difference increases with age |
|  | **Distractors (D)** | Async = Sync, no age variation |
| **Modified Gaze Transition Entropy (mGTE)** | **Distractors (D<->D)** | Async > Sync, no age variation |
|  | **Target (T->D)** | Async > Sync, Async – Sync difference increases with age |
| **Gaze Transition Entropy (GTE)** | **All faces** | Async < Sync, Async-Sync difference increases with age |
| **Rate of Transitions** | **Distractors (D-D)** | Async = Sync, no age variation |
|  | **to/from Target (D->T, T->D)** | Async > Sync, Sync: D->T > T->D |

Table S2. rm-ANOVA of PTLT as calculated Lewkowicz et al. (2021, 2022), where Distractor PTLT corresponds to the average PTLT of all distractors.

|  | **Children** | | | **Adults** | | |
| --- | --- | --- | --- | --- | --- | --- |
|  | *F* | *p* | *ηp²* | *F* | *p* | *ηp²* |
| **Age** | F(4,172)=14.26 | <.001 | .25 | - | - | - |
| **Condition** | F(1,172)=167.98 | <.001 | .49 | F(1,37)=1002.83 | <.001 | 0.96 |
| **Condition x Age** | F(4,172)=9.87 | <.001 | .19 | - | - | - |
| **Stimulus Type** | F(1,172)=235.37 | <.001 | .58 | F(1,37)=855.35 | <.001 | 0.95 |
| **Condition x Stimulus Type** | F(1,172)=167.98 | <.001 | .49 | F(1,37)=1096.24 | <.001 | 0.97 |
| **Stimulus Type x Age** | F(4,172)=14.26 | <.001 | .25 | - | - | - |
| **Condition x Stimulus Type x Age** | F(4,172)=9.87 | <.001 | .19 | - | - | - |

Table S3. rm-ANOVA of PTLT as calculated Lewkowicz et al. (2021, 2022), where Distractor PTLT corresponds to the average PTLT of all distractors¹

|  | **Children** | | | **Adults** | | |
| --- | --- | --- | --- | --- | --- | --- |
|  | *F* | *p* | *ηp²* | *F* | *p* | *ηp²* |
| **Age** | *F*(3,147)=3.24 | .014 | .082 | - | - | - |
| **T Quadrant** | *F*(3,147)=22.57 | < .001 | .336 | *F*(3,45)=35.89 | <. 001 | .50 |
| **Age × T Quadrant** | F(12,147)=1.49 | .196 | .038 | *-* | - | - |

1. For children and adults, rm-ANOVAs included Target Quadrant as a within-subjects factor with data from 143 children and 37 adults. Participants were excluded if they produced no fixation on the target in a Sync trial (see *Participants* in SI Appendix).

Table S4. ANOVA of Target PTLT Scores. ¹

|  | **Children** | | | **Adults** | | |
| --- | --- | --- | --- | --- | --- | --- |
|  | ***F*** | ***p*** | ***ηp²*** | ***F*** | ***p*** | ***ηp²*** |
| **Age** | F(172,4)=14.26 | .001 | .25 |  | - | - |
| **Condition** | F(1,172)=167.98 | <.001 | .49 | F(1,37)=1448.30 | < .001 | .97 |
| **Condition x Age** | F(4,172)=9.87 | <.001 | .19 | - | - | - |
| **T Quadrant** | F(3,170)=25.5 | <.001 | .31 | F(3,111)=10.27 | < .001 | 0.21 |
| **T Quadrant x Age** | F(12,516)=1.32 | .205 | .03 | - | - | - |
| **T Quadrant x Condition** | F(3,170)=3.97 | .009 | .07 | F(3,111)=1.67 | 0.17 | 0.04 |
| **T Quadrant x Condition x Age** | F(12,516)=2.32 | .007 | .05 | - | - | - |

1. As noted in the Methods section of this SI Appendix, our PTLT calculation differed from Lewkowicz et al. (2021; 2022), resulting in a summed target and distractors PTLT of 100%. Given that the distractor PTLT was a complement of target PTLT, we only included the latter measure in the initial ANOVA analyses. Therefore, the within-subjects factors in the rm-ANOVAs were Condition (Sync, Async) and Target Quadrant.

Table S5. ANOVA of Stationary Entropy (STE) across all faces. ¹

|  | **Children** | | | **Adults** | | |
| --- | --- | --- | --- | --- | --- | --- |
|  | *F* | *p* | *ηp²* | ***F*** | ***p*** | ***ηp²*** |
| **Age** | F(4,172) = 18.76 | .049 | .05 | - | - | - |
| **Condition** | F(1,172) = 18.76 | <.001 | .10 | F(1,37) = 410.46 | < .001 | 0.91 |
| **Condition x Age** | F(4,172) = 12.12 | <.001 | .22 | - | - | - |
| **T Quadrant** | F(3,170) = 8.35 | <.001 | .13 | F(3,37) = 16.74 | < .001 | 0.31 |
| **T Quadrant x Condition** | F(3,170) = 2.24, | .085 | .04 | F(3,37) = 15.35 | < .001 | 0.29 |
| **T Quadrant x Age** | F(12,516) = 1.71 | .060 | .038 | - | - | - |
| **T Quadrant x Condition x Age** | F(12,516) = 1.48 | .128 | .033 | - | - | - |

1. To determine whether the differences in PTLT observed across the synchrony conditions and across age reflected changes in fixation dispersion, we computed normalized Shannon’s stationary entropy for fixations on all AOIs. Here, the within-subjects factors in the rm-ANOVAs were Condition, Stimulus Type, and Target Quadrant.

Table S6. ANOVA of Stationary Entropy (STE) on distractors. ¹

|  | **Children** | | | **Adults** | | |
| --- | --- | --- | --- | --- | --- | --- |
|  | ***F*** | ***p*** | ***ηp²*** | ***F*** | ***p*** | ***ηp²*** |
| **Age** | F(4,172)=3.16 | .015 | .07 | - | - | - |
| **Condition** | F(1,172)=2.26 | .134 | .01 | F(1,37)=117.52 | < .001 | 0.76 |
| **Condition x Age** | F(4,172)=3.50 | .009 | .07 | - | - | - |
| **T Quadrant** | F(3,170)=14.9 | <.001 | .21 | F(3,37)=12.20 | < .001 | 0.25 |
| **T Quadrant x Condition** | F(3,170)=.46 | .710 | .01 | F(3,37)=13.94 | < .001 | 0.27 |
| **T Quadrant x Age** | F(12,516)=1.50 | .120 | .01 | - | - | - |
| **T Quadrant x Condition x Age** | F(12,516)=1.45 | .143 | .03 | - | - | - |

1. To determine whether the differences in PTLT observed across the synchrony conditions and across age reflected changes in fixation dispersion, we computed stationary entropy on distractors. Here, the within-subjects factors in the rm-ANOVAs were Condition, Stimulus Type, and Target Quadrant. normalized Shannon’s stationary entropy for fixations on all distractors only.

Table S7. ANOVA of Modified Gaze Transitions (mGTE) from Target to Distractors.¹

|  | **Children** | | | **Adults** | | |
| --- | --- | --- | --- | --- | --- | --- |
|  | ***F*** | ***p*** | ***ηp²*** | ***F*** | ***p*** | ***ηp²*** |
| **Age** | F(4,172)=6.55 | <.001 | .13 | - | - | - |
| **Condition** | F(1,172)=.48 | .488 | .00 | F(1,35)= 123.59 | <0.001 | 0.78 |
| **Condition x Age** | F(4,172)=.27 | .898 | .01 | - | - | - |

1. To characterize the predictability of departure from the target (T->D) and transitions between distractors (D<->D), we computed the modified transitions entropy for these AOIs. Although these measures do not represent the overall transition dynamics, they provide insights into the differences between dynamics of T->D vs. D<->D transitions. The within-subjects factor in this rm-ANOVAs was Condition.

Table S8. ANOVA of Modified Gaze Transitions (mGTE) between Distractors. ¹

|  | **Children** | | | **Adults** | | |
| --- | --- | --- | --- | --- | --- | --- |
|  | ***F*** | ***p*** | ***ηp²*** | ***F*** | ***p*** | ***ηp²*** |
| **Age** | F(4,172)=6.13 | <.001 | .13 | - | - | - |
| **Condition** | F(1,172)=13.40 | <.001 | .07 | F(1,35)= 133.52 | <0.001 | 0.79 |
| **Condition x Age** | F(4,172)=6.26 | <.001 | .13 | - | - | - |

1. To characterize the predictability of departure from the target (T->D) and transitions between distractors (D<->D), we computed the modified transitions entropy for these AOIs. Although these measures do not represent the overall transition dynamics, they provide insights into the differences between dynamics of T->D vs. D<->D transitions. The within-subjects factor in this rm-ANOVAs was Condition.

Table S9. ANOVA of Gaze Transition Entropy (GTE) across all faces. ¹

|  | **Children** | | | **Adults** | | |
| --- | --- | --- | --- | --- | --- | --- |
|  | ***F*** | ***p*** | ***ηp²*** | ***F*** | ***p*** | ***ηp²*** |
| **Age** | F(4,172)=3.63 | .007 | .08 | - | - | - |
| **Condition** | F(1,172)=3.77 | .054 | .021 | F(1,35)=177.84 | <.001 | .84 |
| **Condition x Age** | F(4,172=.53 | 0.716 | .01 | - | - | - |

1. To characterize the overall dynamics underlying the stationary distribution of gaze (Target and Distractor PTLT as well as Stationary Entropy), we computed the transitions entropy for transitions between all faces, weighted by dwell-time on each face. Condition was the within-subjects factor in the rm-ANOVAs.

Table S10. ANOVA of Rate of Transitions between Distractors over Total Distractors Fixation Time.¹

|  | **Children** | | | **Adults** | | |
| --- | --- | --- | --- | --- | --- | --- |
|  | ***F*** | ***p*** | ***ηp²*** | ***F*** | ***p*** | ***ηp²*** |
| **Age** | F(4,167)=2.38 | .60 | .02 | - | - | - |
| **Condition** | F(1,167)=4.96 | .027 | .028 | F(1,35)=29.41 | <.001 | .456 |
| **Condition x Age** | F(4,167)=.69 | 0.596 | .02 | - | - | - |

1. To assess the likelihood of transitioning between distractors, we divided the number of such transitions by the summed duration of fixations on distractors. The rm-ANOVAs included only Condition as within-subjects factor.

Table S11. ANOVA of Rate of Transitions between Target and Distractors Transitions over Total Target Fixation Time.¹

|  | **Children** | | | **Adults** | | |
| --- | --- | --- | --- | --- | --- | --- |
|  | ***F*** | ***p*** | ***ηp²*** | ***F*** | ***p*** | ***ηp²*** |
| **Age** | F(4,172)=.96 | .430 | .02 | - | - | - |
| **Condition** | F(1,172)=57.20 | <.001 | .25 | F(1,33)= 390.76 | <.001 | .92 |
| **Condition x Age** | F(4,172)=10.30 | .201 | .03 | - | - | - |
| **Direction** | F(1,172)=.20 | .081 | .02 | F(1,33)=3.17 | .08 | .09 |
| **Direction x Condition** | F(1,172)=2.88 | .091 | .02 | F(1,33)=2.51 | .12 | .07 |
| **Direction x Age** | F(4,172)=4.03 | .004 | .09 | - | - | - |
| **Direction x Condition x Age** | F(4,172)=.16 | .958 | .00 | - | - | - |

1. To assess the likelihood of transitioning between distractors, we divided the number of such transitions by the summed duration of fixations on distractors. The rm-ANOVAs included Condition as the only within-subjects factor.

**Movie S1 (separate file).** Actor: Alyssa, Condition: Sync, Target Quadrant: Left Bottom, Utterance: "We can...".

**Movie S2 (separate file).** Actor: Alyssa, Condition: Sync, Target Quadrant: Left Bottom, Utterance: "Good morning.".

**Movie S3 (separate file).** Actor: Alyssa, Condition: Sync, Target Quadrant: Left Top, Utterance: "Good morning.".

**Movie S4 (separate file).** Actor: Alyssa, Condition: Sync, Target Quadrant: Left Top, Utterance: "We can...".

**Movie S5 (separate file).** Actor: Alyssa, Condition: Sync, Target Quadrant: Right Bottom, Utterance: "Good morning.".

**Movie S6 (separate file).** Actor: Alyssa, Condition: Sync, Target Quadrant: Right Bottom, Utterance: "We can...".

**Movie S7 (separate file).** Actor: Alyssa, Condition: Sync, Target Quadrant: Right Top, Utterance: "Good morning.".

**Movie S8 (separate file).** Actor: Alyssa, Condition: Sync, Target Quadrant: Right Top, Utterance: "We can...".

**Movie S9 (separate file).** Actor: Alyssa, Condition: Async, Target Quadrant: Left Bottom, Utterance: "Good morning.".

**Movie S10 (separate file).** Actor: Alyssa, Condition: Async, Target Quadrant: Left Bottom, Utterance: "We can...".

**Movie S11 (separate file).** Actor: Alyssa, Condition: Async, Target Quadrant: Left Top, Utterance: "Good morning.".

**Movie S12 (separate file).** Actor: Alyssa, Condition: Async, Target Quadrant: Left Top, Utterance: "We can...".

**Movie S13 (separate file).** Actor: Alyssa, Condition: Async, Target Quadrant: Right Bottom, Utterance: "Good morning.".

**Movie S14 (separate file).** Actor: Alyssa, Condition: Async, Target Quadrant: Right Bottom, Utterance: "We can...".

**Movie S15 (separate file).** Actor: Alyssa, Condition: Async, Target Quadrant: Right Top, Utterance: "Good morning.".

**Movie S16 (separate file).** Actor: Alyssa, Condition: Async, Target Quadrant: Right Top, Utterance: "We can...".

**Movie S17 (separate file).** Actor: Megan, Condition: Sync, Target Quadrant: Left Bottom, Utterance: "But...".

**Movie S18 (separate file).** Actor: Megan, Condition: Sync, Target Quadrant: Left Bottom, Utterance: "They like to ice...".

**Movie S19 (separate file).** Actor: Megan, Condition: Sync, Target Quadrant: Left Top, Utterance: "But...".

**Movie S20 (separate file).** Actor: Megan, Condition: Sync, Target Quadrant: Left Top, Utterance: "They like to ice...".

**Movie S21 (separate file).** Actor: Megan, Condition: Sync, Target Quadrant: Right Bottom, Utterance: "But...".

**Movie S22 (separate file).** Actor: Megan, Condition: Sync, Target Quadrant: Right Bottom, Utterance: "They like to ice...".

**Movie S23 (separate file).** Actor: Megan, Condition: Sync, Target Quadrant: Right Top, Utterance: "But...".

**Movie S24 (separate file).** Actor: Megan, Condition: Sync, Target Quadrant: Right Top, Utterance: "They like to ice...".

**Movie S25 (separate file).** Actor: Megan, Condition: Async, Target Quadrant: Left Bottom, Utterance: "But...".

**Movie S26 (separate file).** Actor: Megan, Condition: Async, Target Quadrant: Left Bottom, Utterance: "They like to ice...".

**Movie S27 (separate file).** Actor: Megan, Condition: Async, Target Quadrant: Left Top, Utterance: "But...".

**Movie S28 (separate file).** Actor: Megan, Condition: Async, Target Quadrant: Left Top, Utterance: "They like to ice...".

**Movie S29 (separate file).** Actor: Megan, Condition: Async, Target Quadrant: Right Bottom, Utterance: "But...".

**Movie S30 (separate file).** Actor: Megan, Condition: Async, Target Quadrant: Right Bottom, Utterance: "They like to ice...".

**Movie S31 (separate file).** Actor: Megan, Condition: Async, Target Quadrant: Right Top, Utterance: "But...".

**Movie S32 (separate file).** Actor: Megan, Condition: Async, Target Quadrant: Right Top, Utterance: "They like to ice...".

**SI References**

1. C. Chandrasekaran, A. Trubanova, S. Stillittano, A. Caplier, A. A. Ghazanfar, The natural statistics of audiovisual speech. PLoS Comput Biol 5, e1000436 (2009).
2. B. Shiferaw, L. Downey, D. Crewther, A review of gaze entropy as a measure of visual scanning efficiency. Neurosci Biobehav Rev 96, 353–366 (2019).

1. The four utterances consisted of the following: (1) "But your favorite will be the elephants. They’re big and gray and have large floppy ears. Maybe we’ll see a baby elephant too? What do you think about that? If not, we could go to story time at the library. All your friends will be there"; (2) "They like to ice skate, right? But, before we can go anywhere, what do we have to do? Change your clothes and eat breakfast, of course. It’s cold outside, so you need to wear a sweater. How about the green one with the duck? For breakfast, you can have oatmeal with blueberries."; (3) "Good morning, get up, come on now. If you get up right away, we’ll have an hour to play in the house. I love these long mornings, don’t you. I wish they could last all day."; (4) "We can hang around all day Saturday. Except, of course, for the party. Are you going to help me fix up the house? Are you? We need to buy flowers, prepare the food, vacuum the house, dust."
